# MyoD-Cre driven alterations in K-Ras and p53 lead to a mouse model with histological and molecular characteristics of human rhabdomyosarcoma with direct translational applications

**DOI:** 10.1101/2021.06.15.448607

**Authors:** Kengo Nakahata, Brian W. Simons, Enrico Pozzo, Ryan Shuck, Lyazat Kurenbekova, Cristian Coarfa, Tajhal D. Patel, Jason T. Yustein

## Abstract

Rhabdomyosarcoma (RMS) is the most common soft tissue sarcoma in children, with overall long-term survival rates of about 65-70%. Thus, additional molecular insights and representative models are critical for further identifying and evaluating new treatment modalities. Using MyoD-Cre mediated introduction of mutant K-Ras^G12D^ and perturbations in p53 we have developed a novel genetically engineered mouse model (GEMM) for RMS. Specifically, we directly crossed mice expressing MyoD promoter-regulated Cre-recombinase with germline p53^Flox^ or Lox-Stop-Lox (LSL) knock-in alleles expressing oncogenic p53^R172H^ and/or K-Ras^G12D^ mutants. The anatomic sites of primary RMS development observed in these mice recapitulated human disease, with the most frequent sites of tumor growth seen in the head, neck, extremities, and abdomen.

We have confirmed RMS histology and diagnosis through hematoxylin and eosin (H&E) staining, as well as positive immunohistochemistry (IHC) staining for desmin, myogenin, and phosphotungstic acid hematoxylin (PTAH). We established cell lines from several of the GEMM tumors with the ability to engraft and develop tumors in immunocompetent mice with similar histological and staining features as the primary tumors. Furthermore, injection of syngeneic RMS lines via tail vein had high metastatic potential to the lungs. Transcriptomic analyses of p53^R172H^/K-Ras^G12D^ GEMM-derived tumors showed evidence of high molecular homology with human RMS. Specifically, we noted alterations in gene ontologies including immune response, metabolism and mRNA processing. Finally, pre-clinical use of these murine RMS lines demonstrated similar therapeutic responsiveness to relevant chemotherapy and targeted therapies as human cell line models.

**Summary Statement:** We have developed a new conditional genetically engineered mouse model of rhabdomyosarcoma (RMS) with homologous molecular signature to human RMS that provides valuable pre-clinical models for evaluating novel therapies.

## Introduction

Rhabdomyosarcoma (RMS) is the most common soft tissue sarcoma in children. Despite improvements in patient survival rates over the past few decades, over one-third of RMS patients still succumb to the disease. Therefore, identification of additional effective therapeutic regimens is crucial towards improving the outcome for these patients. RMS can be categorized histologically into alveolar RMS (ARMS) and embryonal RMS (ERMS) (Yohe et al., 2019). The alveolar subtype has a distinct alveolar architecture with aggregates of small round undifferentiated cells separated by dense hyalinized fibrous septa. At the center of the tumor, the clusters are arranged loosely and therefore they look like an alveolar. The embryonal subtype resembles microscopically the various stages of muscle development from poorly differentiated round tumor cells to well-differentiated, with cross-striations like rhabdomyoblasts (Wang, 2012).

Furthermore, ARMS typically harbor a chromosomal translocation, PAX3 or 7 /FOXO1 (FKHR), rendering ARMS to be grouped as fusion positive, and ERMS as fusion negative. Although ARMS rarely involves the inactivation or mutations of p53, ERMS has a relatively high frequency of mutations or aberrations in the p53/MDM2 axis (Felix et al., 1992). In addition, metastases in ERMS show high levels of p53 mutations (Leuschner et al., 2003). The K-Ras oncogene has been implicated in various cancers, with up to one-third of RMS cases involving the activation of one of the three Ras isoforms, including K-Ras (Ku et al., 2017; Stratton et al., 1989; Wilke et al., 1993).

Although rapid progress has been made in the field of RMS, most researches are based on xenotranslantation using human cell lines into immunodeficient models. These xenograft models do not recapitulate the complete tumor biology, including the tumor immune microenvironment. Thus, there is a significant need to develop murine RMS models that can develop and progress in immunocompetent model systems.

In this study, we have developed and characterized a conditional p53 and/or K-Ras fusion-negative RMS genetically engineered mouse model (GEMM). We also performed histologic and genetic analysis of GEMM tumors, and found that these tumors had similar phenotypic and molecular features to human RMS. Based on the homologous histological and molecular traits to RMS we believe that this mouse model, and the derived cell lines could signficantly contribute to our understanding of the molecular pathogenesis of RMS and provide valuable immunocompetent models for investigating novel, innovative therapeutic strategies for treating RMS.

## Results

### Generating and characterizing non-metastatic and metastatic rhabdomyosarcoma mouse models

To develop a RMS genetically engineered mouse model, we crossed MyoD-Cre mice with germline Lox-Stop-Lox (LSL) p53^R172H^ or K-Ras^G12D^ alleles. Four distinct MyoD-Cre transgenic genotypes were monitored for tumor incidence: K-Ras-WT mice with floxed p53 allele/wt p53 allele or LSL-p53R172H allele/wt p53 allele (Cre+ F/+ and Cre+R/+) and K-Ras-mutated mice with floxed p53 allele/wt p53 allele or LSL-p53R172H allele/wt p53 allele (Cre+K-Ras^G12D^;F/+ and Cre+K-Ras^G12D^;R/+) (Figure 1A).

**Figure 1.**
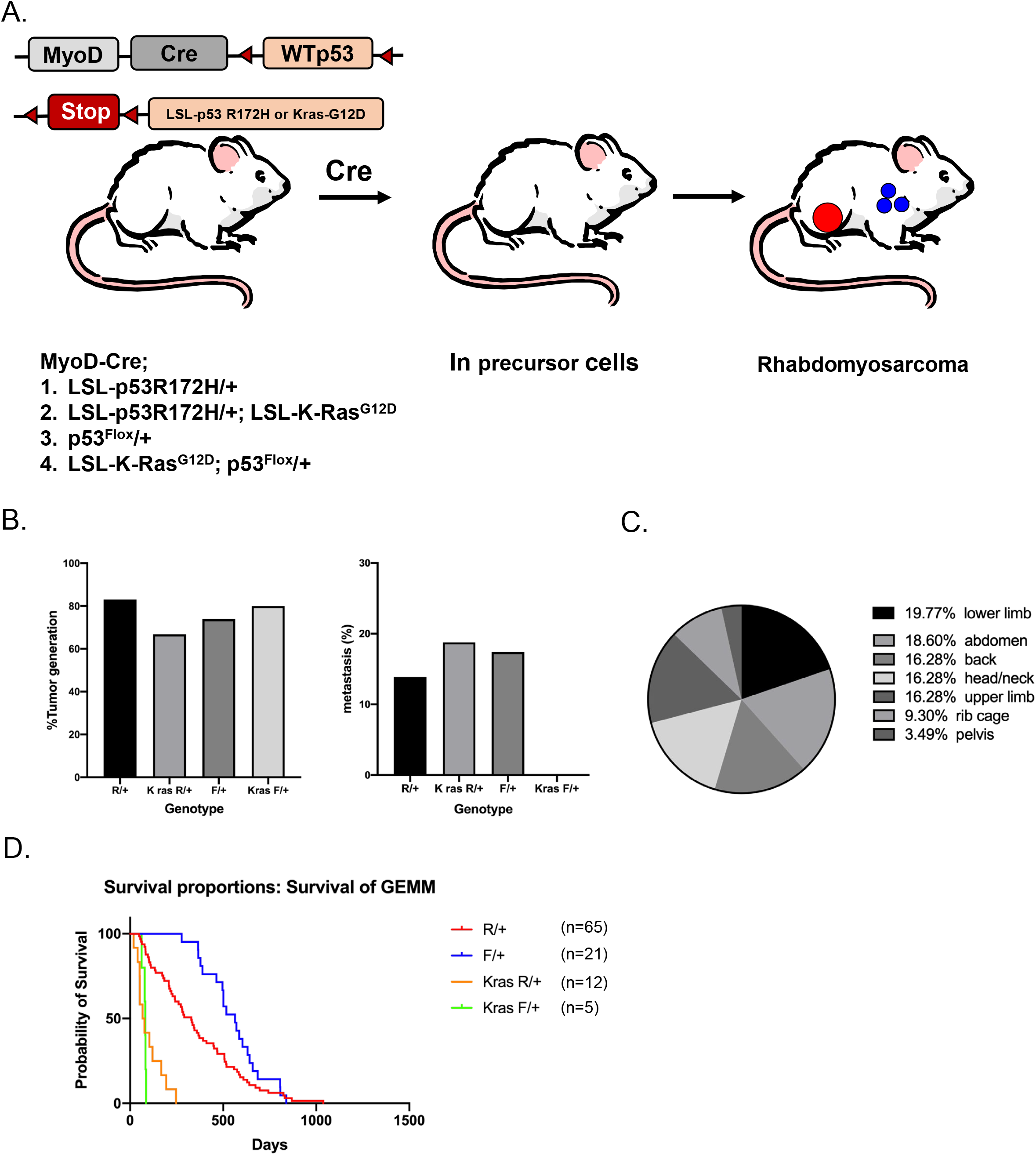
Design and characterization of a novel genetically engineered mouse model of metastatic rhabdomyosarcoma. (A)Schematic representing rhabdomyosarcoma-susceptible mice. All mice expressed germline MyoD-Cre allele in combination with (1) heterozygous deletion of p53 alone (F/+), (2) LSL-p53R172H gain-of-function mutant p53 (R/+), (3) LSL-K-RasG12D gain-of-function K-Ras mutant in combination with heterozygous deletion of p53 (KRas;F/+) or (4) LSL-K-RasG12D gain-of-function K-Ras mutant in combination with LSL-p53R172H gain-of-function mutant p53 (KRas; R/+). (B) Tumor incidence frequency (%) for each genotype and frequency (%) of metastatic rhabdomyosarcomas varies according to p53 genotype and K-Ras mutation. ‘F’=floxed p53, ‘R’=LSL-p53R172H, and ‘+’=WTp53 allele. (C) Anatomical distribution of primary tumor location for rhabdomyosarcoma of the different K-Ras and p53 genotypes. (D) Kaplan-Meier plots for each of four different genotypes of mice.

During necropsy, we assessed for metastatic lesions in all of the major organs, noting lungs to be the most frequent site of metastasis, which we noted ranged between 10-20% of the animals, while there seemed to be no significant difference in incidence of tumor generation between each group (Figure 1B). The sites of RMS development varied, with the lower limbs showing the highest frequency of lesions (19.8%), followed by abdomen (18.2%), back, head/neck, and upper limbs (15.9% each) (Figure 1C). These results suggest that murine RMS could develop in multiple anatomical locations that are comparable to those seen in human patients.

We examined the overall survival of these genetically engineered mice, comparing the survival rates of K-Ras-mutated with K-Ras-WT mice (Figure 1D). We noted that K-Ras-WT mice (Cre+ F/+ and Cre+R/+) lived significantly longer than K-Ras-mutated mice (Cre+K-Ras^G12D^;F/+ and Cre+K-Ras^G12D^;R/+, *p*<0.0001). There were no statistically significant differences between the K-Ras^G12D^ cohorts (*p*=0.58).

### Immunohistochemistry analysis of GEMM tumors

Following the evaluation of tumor incidence, primary tumor development and metastatic potential in this murine model, we next sought to characterize the tumors histologically. Tumor morphology was evaluated by a veterinary pathologist with experience in mouse sarcoma models, and skeletal muscle differentiation was evaluated using PTAH histochemical staining and IHC for desmin and myogenin, markers of muscle and skeletal muscle differentiation respectively (Fig 2A-C). Primary tumors were moderate to poorly differentiated sarcomas, typically with round to spindloid cells with eosinophilic cytoplasm and pleomorphic nuclei. Most tumors had identifiable elongated “strap” cells or cells with visible cross striations that are specific for rhabdomyosarcoma, but a proportion of primary tumors were undifferentiated or anaplastic sarcomas. Most tumors showed RMS histology and moderate to high desmin and myogenin expression by immunohistochemistry.

**Figure 2.**
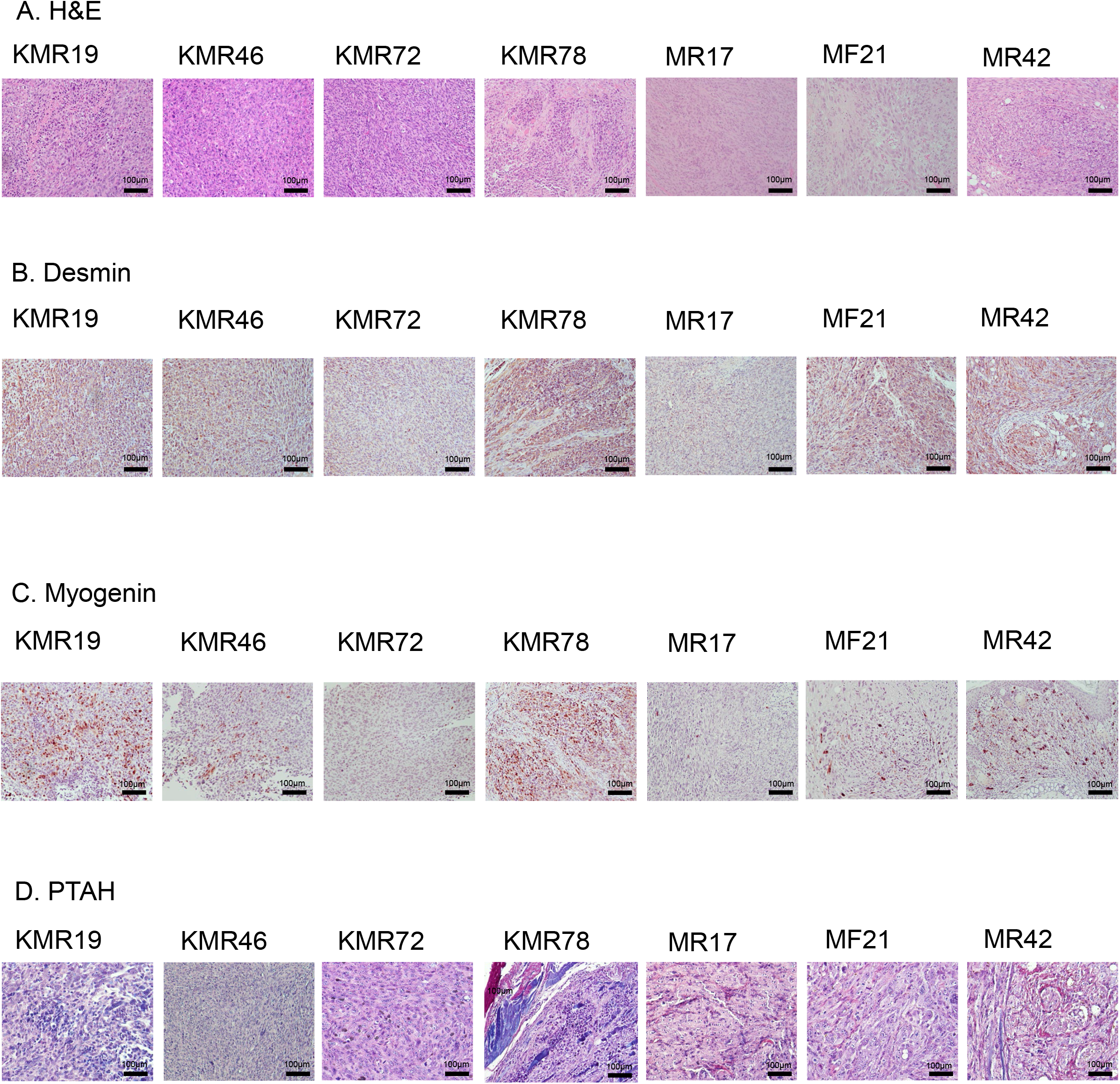
Tumor histopathological analysis of GEMM RMS tumors. (A) Representative H&E staining for GEMM tumors. (B) Desmin immunohistochemistry (IHC) staining of GEMM tumors. They were analyzed by hematoxylin and Desmin. (C) Myogenin IHC staining of GEMM tumors. They were analyzed by hematoxylin and myogenin. (C) Phosphotungstic acid hematoxylin (PTAH) staining of GEMM tumors. Muscle is stained blue black to blue purple, connective tissue is orange pink to brownish red, fibrin and neuroglia stain deep blue, and coarse elastic fibers show as purple. Scale bars correspond 100 μm.

### Molecular characterization

We performed RNA-sequencing from primary GEMM tumors to examine whether gene expression in tumors from our model overlapped with gene expression changes seen in human RMS. For our studies, we used K-Ras^G12D^ tumors, K-Ras^WT^ tumors and normal gastrocnemius muscle tissue from three non-tumor bearing mice as controls. We generated gene signatures of at least 1.5 fold change and an FDR ≤ 0.05 for K-Ras ^G12D^ or K-Ras ^WT^ compared to normal gastrocnemius muscle tissue, as well as human RMS (either fusion negative or fusion positive) to normal muscle tissue. To determine the extent of overlap between the differentially expressed genes (DEGs) from the K-Ras ^G12D^ and K-ras^WT^ tumors with human RMS, we compared the murine DEGs to the human FP-RMS and FN-RMS gene signatures. Of the 7900 FN-RMS DEGs, 30% (2391 genes) and 45% (3585 genes) overlapped with our K-Ras^WT^ and K-Ras ^G12D^ respectively (Figure 3A). Similarly, the 8247 FP-RMS DEGs had a 28% (2333 genes) overlap with K-Ras^WT^ and 43% (3547 genes) overlap with K-Ras^G12D^ (Figure 3A).

**Figure 3.**
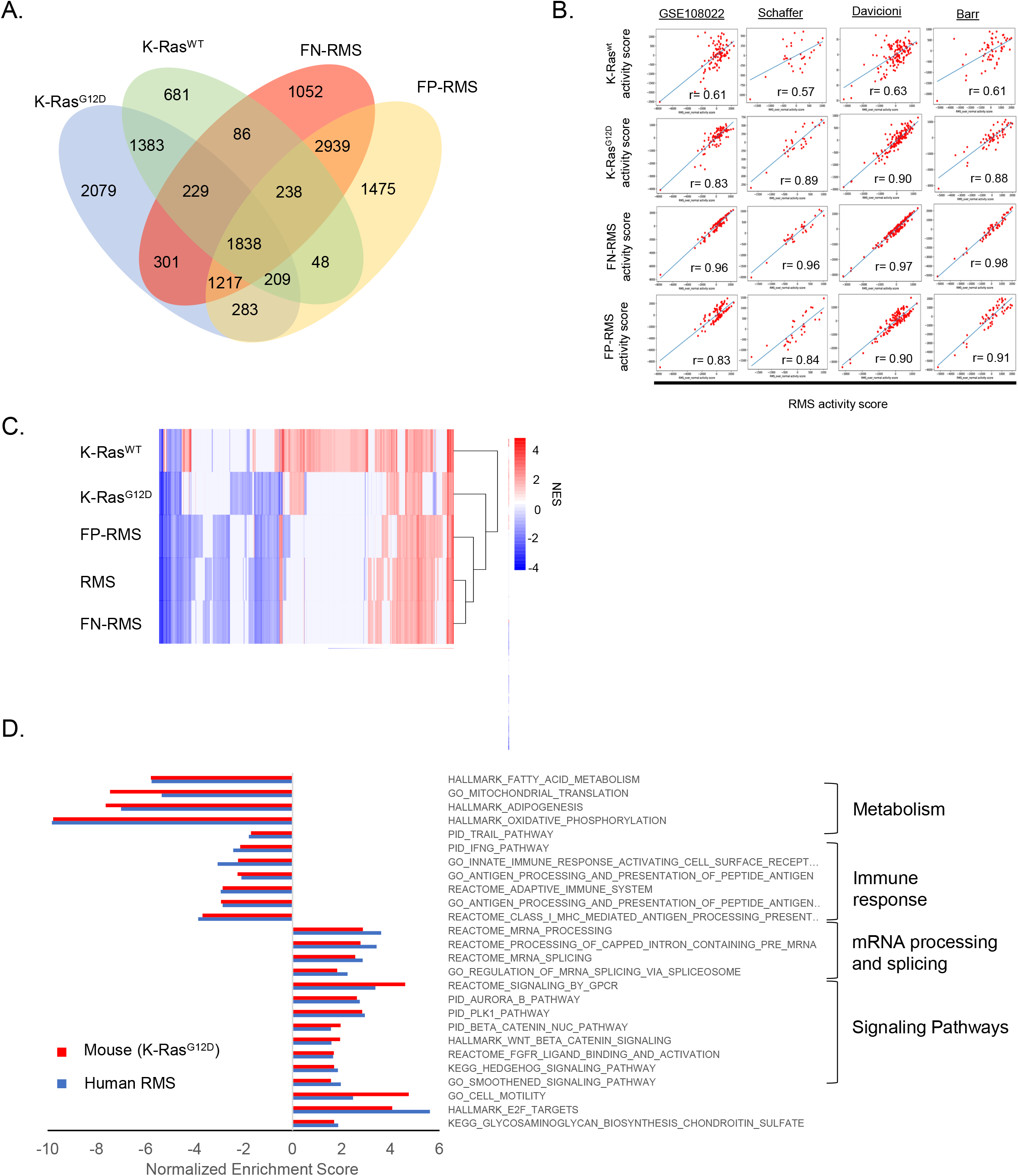
Molecular characterization of RMS GEMM tumors. (A) Overlap genes with human fusion positive and negative gene signatures. ‘NES’= normalized enrichment score. (B) Pearson correlation coefficients using GSE108022, Schaffer, Davicioni and Barr. (C) Gene Set Enrichment Analysis (GSEA) using the mouse and human DEG signatures. ‘NES’= normalized enrichment score. (D) Normalized Enrichment Scores compared between the K-Ras^G12D^ and human RMS. Scale bars correspond 100 μm.

We further asked whether the overlapping gene changes were consistent across multiple human RMS datasets. We performed signature correlation between our human RMS gene signature derived from GSE108022 and murine K-Ras gene signatures in other human RMS cohorts. While both the K-Ras ^G12D^ and K-Ras ^WT^ signatures were significantly correlated with our human RMS signature in all four cohorts analyzed, the K-Ras ^G12D^ signature consistently had much higher Pearson correlation coefficients, similar to fusion-negative RMS (FN-RMS) and fusion-positive RMS (FP-RMS), compared to K-Ras ^WT^ (Figure 3B). A previous report by Blum et al. reported that a mouse model of MyoD-expressing cells with *ex vivo* activation of oncogenic K-Ras and p53 deletion led to the formation of Undifferentiated Pleiomorphic Sarcomas (UPS) as opposed to RMS (Blum et al., 2013). We therefore also examined the correlation of our mouse model signatures to UPS DEGs and found the K-Ras^WT^ signature had a higher correlation coefficient (0.97) with the UPS gene signature compared to K-Ras^G12D^ (0.89), which had comparable correlation coefficients to UPS with human FN- and FP-RMS (Supplemental Fig. 1). Taken together these data indicate that the K-Ras ^G12D^ GEMM model more closely mimics the gene expression changes seen in human RMS across multiple datasets when compared to the K-Ras^WT^ model, which has gene expression changes more closely related to those seen in human UPS.

To better examine whether the K-Ras^G12D^ mouse model captures human RMS pathway changes we ran Gene Set Enrichment Analysis (GSEA) using the mouse and human DEG signatures. Interestingly, based on unbiased hierarchical clustering using the resulting normalized enrichment scores (NES) the K-Ras ^G12D^ clustered closer to human RMS than the K-Ras ^WT^ (Figure 3C) suggesting that the K-Ras ^G12D^ model is more representative of the human disease than K-Ras ^WT^. More importantly, both the K-Ras^G12D^ and human RMS showed suppression of gene sets related to metabolism and immune response while mRNA processing, mRNA splicing, and Sonic Hedgehog pathway, among other oncogenic signaling cascades, were enhanced (Figure 3D), thus indicating a recapitulation of the human phenotype on a molecular level.

Finally, we performed Connectivity Map analysis (Subramanian et al., 2017) with the top 150 upregulated K-Ras^G12D^ genes and identified multiple drugs with high reversal scores that have known effects in RMS (Table 1). Vincristine, in particular is an approved treatment for RMS and has been used in conjunction with irinotecan in clinical trials. FGFR and MEK inhibitors have also shown promise in laboratory studies, further supporting our overall analysis that our GEMM model has homologous molecular signatures to the human disease.

**Table 1.** Connectivity Map analysis

| Score | ID | Name | Description |
| --- | --- | --- | --- |
| -99.72 | BRD-K28428262 | brivanib | FGFR inhibitor |
| -99.05 | BRD-K91696562 | orantinib | FGFR inhibitor |
| -97.11 | BRD-K49865102 | PD-0325901 | MEK inhibitor |
| -95.75 | BRD-K89014967 | AS-703026 | MEK inhibitor |
| -95.64 | BRD-K82109576 | vincristine | Tubulin inhibitor |
| -93.29 | BRD-K08547377 | irinotecan | Topoisomerase inhibitor |

### RMS phenotype is maintained in cell lines and orthotopic allografts

After isolation of GEMM tumors, as outlined in our methods, we developed primary cell lines from several GEMM tumors. K-Ras ^G12D^ cell lines KMR19 (R/+), KMR46 (F/+) and KMR78 (R/+), were injected into the gastrocnemius of C57BL/6J mice (Fig. 4A). Resulting orthotopic tumors were resected, stained for desmin and myogenin and examined histologically, which demonstrated their resemblance to their primary GEMM tumors (Fig. 4B&C). In addition, for comparative purposes, we also stained orthotopic human tumor models for desmin and myogenin derived from RMS cell lines RD and Rh30 as well as from an institutional RMS patient-derived xenograft (PDX), TCCC-RMS40 (PDX40) (Fig. 4B&C). The staining patterns observed were again consistent with RMS, showing a high correlation between GEMM and human tumors. Of note, we attempted to develop syngeneic orthotopic tumors from MR17, MF21 and MR42 in C57BL/6 mice, but after several months (2-4 months) no tumors were evident upon necropsy of the animals.

**Figure 4.**
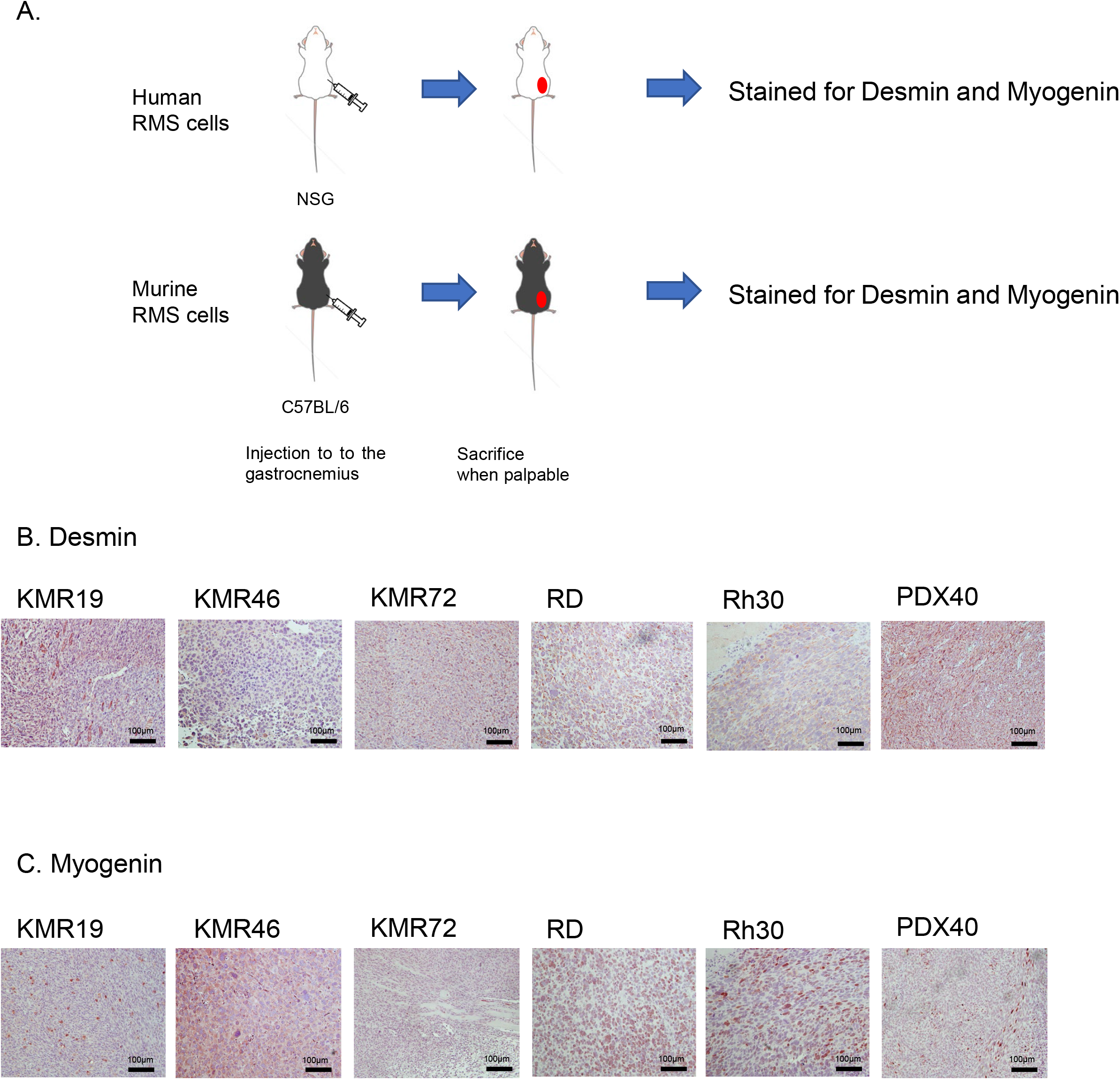
Histopathological analysis of orthotopic transplantation into immunocompetent mice. (A)The schema of orthotopic RMS tumor studies. The cell lines (RD, Rh30, KMR19, KMR46 and KMR72) and PDX40 were injected to the gastrocnemius. The tumors were resected when they were palpable. (B) Desmin immunohistochemistry (IHC) staining of orthotopic RMS tumors. They were analyzed by hematoxylin and Desmin. (C) Myogenin immunohistochemistry (IHC) staining of orthotopic RMS tumors. They were analyzed by hematoxylin and Myogenin. Scale bars correspond 100 μm.

### Pathology of metastatic tumors from murine cell lines

To assess the metastatic potential of these murine cell lines, KMR cell lines were injected via tail vein into C57BL/6J mice. One week following injection, mice were sacrificed and examined for metastatic disease. We observed that all mice showed evidence of macroscopic lung metastasis, which was further analyzed by H&E staining (Figure 5). No metastases were observed in other organs.

**Figure 5.**
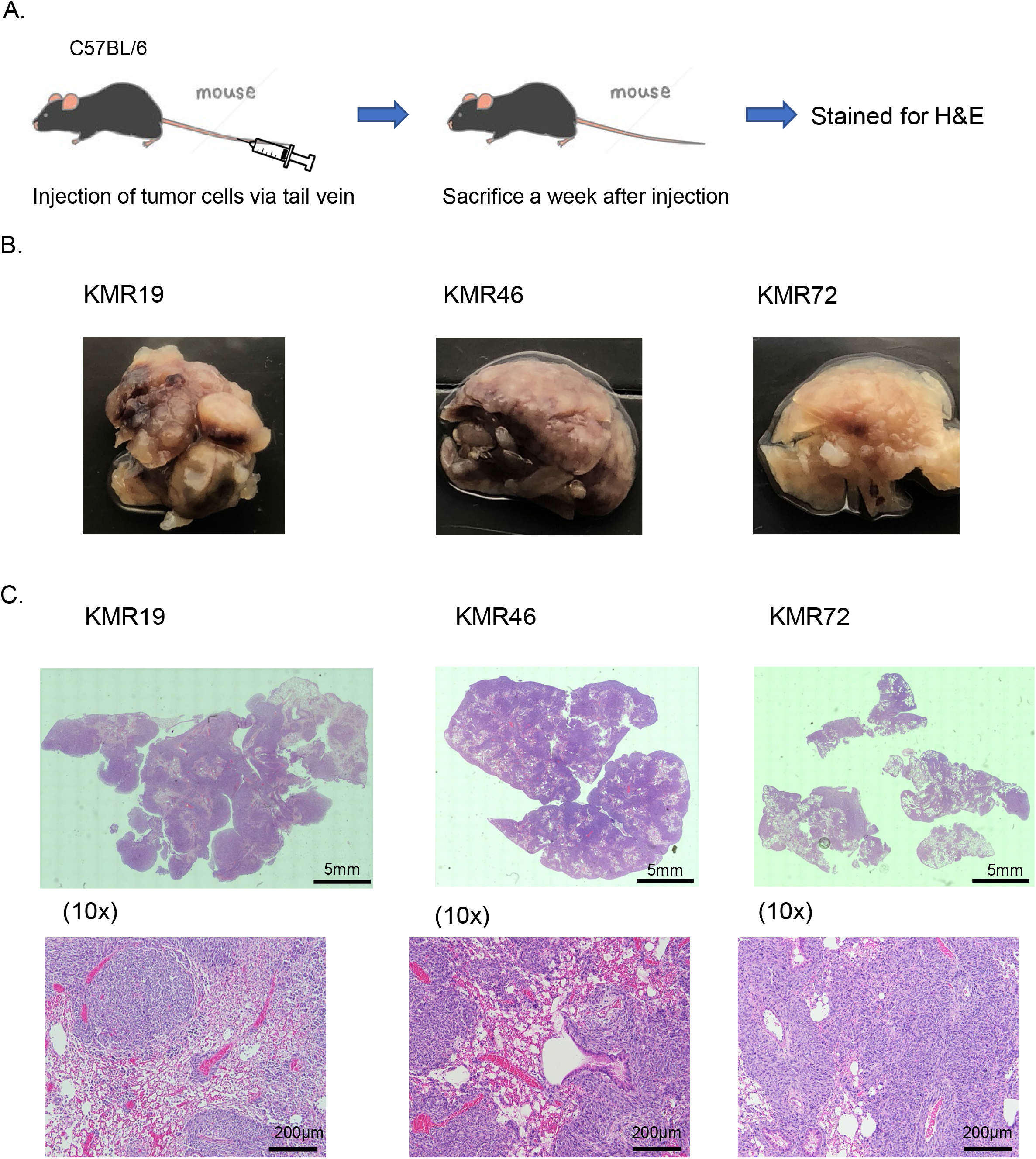
In vivo metastatic potential of GEMM-derived RMS cell lines in immunocompetent mice. After 1 week post tail vein injection, animals were sacrificed and (A) brightfield images of whole lungs using tile scans (as described in materials and methods) and (B) H&E staining of lung lesions was performed for lung sections demonstrating large number of metastatic lesions present. Whole lung and 10x magnification are shown. Scale bars correspond 5 mm (whole lung) and 200 μm (10x).

### Cross-species comparison of RMS cell line models for sensitivity to anti-cancer agents

Based on our Connectivity Map results (Table 1) we sought to determine if anticancer drugs commonly used to treat human RMS were effective treatments for GEMM-derived RMS disease. We performed cytotoxicity assays to assess cell viability in tumor-derived cell lines treated with vincristine and actinomycin D. We evaluated chemosensitivity for three murine cell lines, KMR19, KMR46 and KMR78, which were established from K-Ras ^G12D^ murine tumors, as well as human RMS cell lines, RD and Rh30. Besides the aforementioned chemotherapy, we investigated targeted anti-cancer agents, including a Wee1 inhibitor, MK-1775, and the MEK inhibitor, Trametinib. Our rationale for investigating these latter targets include prior reports demonstrating that Wee1 is a viable therapeutic target, in combination with chemotherapy, in pre-clinical models of RMS (Stewart et al., 2017) (Yohe et al., 2019), while MEK inhibition has been shown to have anti-tumor efficacy in pre-clinical FN-RMS models (Marampon et al., 2006; Yohe et al., 2018). Our findings revealed that all chemotherapeutic agents were effective in treating the murine cell lines, with highly comparable IC_50_ values between our murine RMS lines and the human RMS cell lines (Table 2 and Figure 6). Overall, these *in vitro* pre-clinical studies provide evidence that our RMS GEMM-derived models are extremely valuable resources to evaluate and test novel therapeutic regimens for RMS.

**Table 2.** Cell viability assay on rhabdomyosarcoma cell lines treated with chemotherapeutic agents

|  | IC<sub>50</sub> (95% CI) nM |  |  |  |
| --- | --- | --- | --- | --- |
|  | Vincristine | Actinomycin D | MK1775 | Trametinib |
| RD (human) | 7.58 (6.56-9.00) | 1.46 (0.67-2.23) | 211.8 (150.0-290.6) | 13.32 (3.82-20.69) |
| Rh30 (human) | 2.52 (2.16-3.00) | 1.87 (1.29-2.31) | 135.9 (113.1-172.1) | 149.8 (121.1-187.9) |
| KMR19 (mouse) | 19.29 (14.90-24.56) | 1.65 (1.50-1.80) | 136.2 (112.1-160.9) | 34.03 (7.80-52.31) |
| KMR46 (mouse) | 35.19 (22.51-405.3) | 3.424 (2.915-4.013) | 320.9 (221.6-503.0) | 28.49 (6.54-43.66) |
| KMR78 (mouse) | 15.49 (10.98-23.86) | 1.23 (1.05-1.48) | 35.47 (31.64-38.99) | 34.69 (25.32-44.68) |

**Figure 6.**
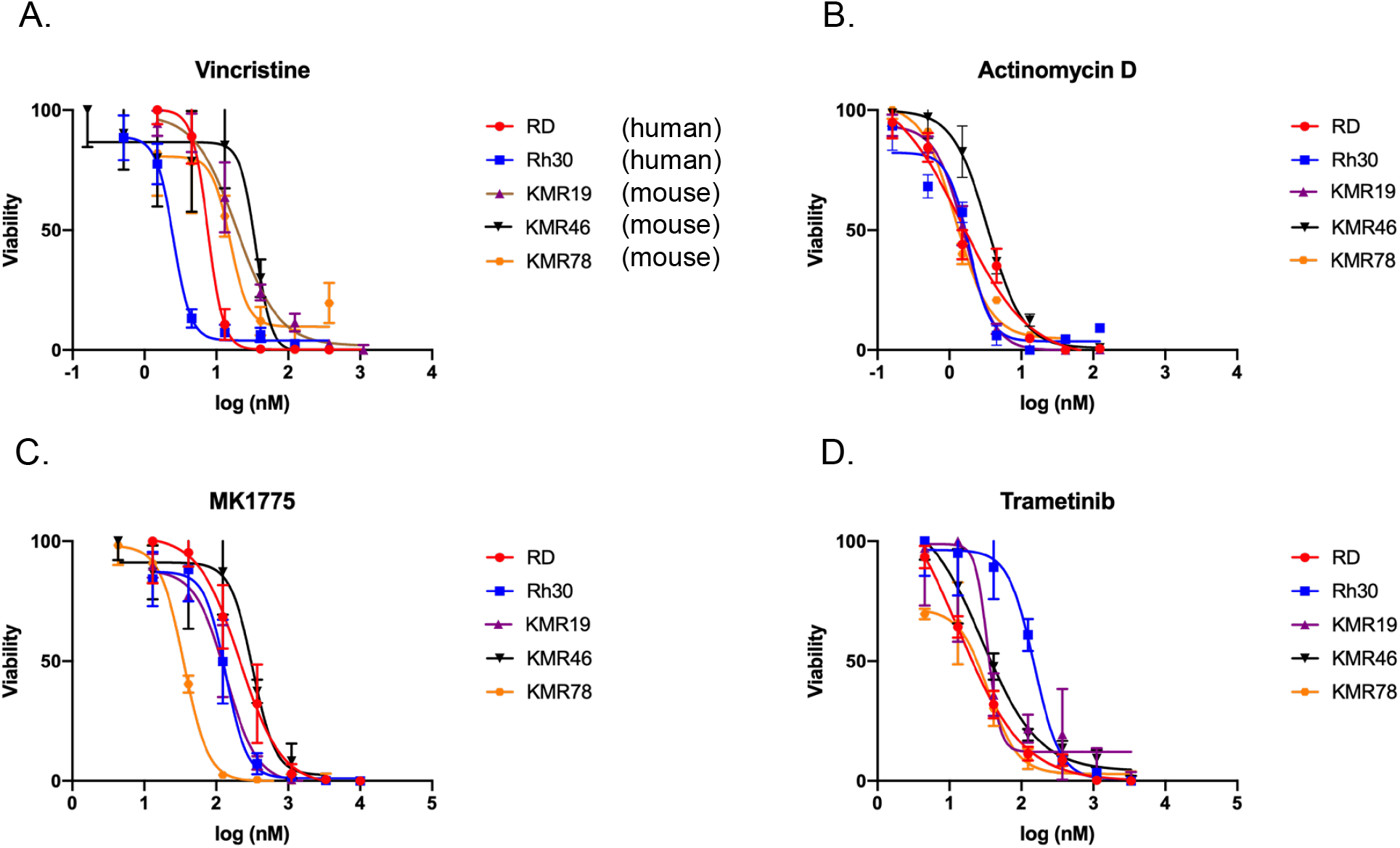
In vitro assessment of therapeutic responsiveness of murine RMS cell lines. Comparison analysis of cytotoxicity for human (RD and Rh30) and murine (KMR19, KMR46 and KMR78) cell lines for (A) Vincristine, (B) Actinomycin D, (C) MK1775, and (D) Trametinib . The viability was measured by CCK-8 (Cell Counting Kit-8) assay and the IC_50_ of each agent is shown in Table 2. Data presented as mean ± SD.

## Discussion

Due to a limited availability of pediatric patient tumor samples, genetically-engineered mouse models and derived cell lines offer valuable resources for gaining insights into the molecular pathogenesis of tumor development and progression as well as immunocompetent pre-clinical models for evaluating the efficacy of targeted mono- and combination therapies, including immune modulatory agents.

Through *in utero* conditional perturbations of K-Ras and p53 in skeletal muscle precursor cells we have established a novel genetically engineered mouse model of rhabdomyosarcoma that exhibits high penetrance and development of histologically analogous primary tumors in homologous anatomical sites seen in RMS patients. Cross-species molecular analysis of our GEMM with human RMS transcriptomic data revealed significant overlap among key human RMS gene ontology and signaling pathways that emphasize critical molecular events driving RMS development. Furthermore, a subpopulation of our GEMM demonstrated evidence of metastatic disease, thus providing additional resources for analysis of the pathogenesis of metastatic disease. Molecular analysis of the metastatic lesions are actively ongoing and beyond the scope of this present study. We did note some limitations to these studies. First, simultaneous occurrence, or multifocal disease was found in some mice, which is rare in RMS. Additionally, the survival times of K-Ras-mutated mice were shorter than non-mutated mice, which makes it difficult to examine the effects of K-Ras on tumor generation and growth.

While there have been previous fusion-negative RMS murine models reported (Hettmer et al., 2011; Rubin et al., 2011) (McKinnon et al., 2015), our model differs from those through *in utero* genetic perturbations that drive endogenous and spontaneous development of RMS. In these models we were able to get spontaneous metastatic tumors in the lung without venous injection and isolate syngenic K-ras mutated cell lines which helps us to investigate murine RMS. Interestingly, prior reports of various skeletal precursor conditional RMS models have demonstrated discrepancy between PAX7 and MyoD-driven recombinase. According to the report by Blum et al., they used PAX7-CreER and MyoD-CreER mice to transform Pax7^+^ and MyoD^+^ myogenic progenitors by expressing oncogenic K-Ras^G12D^ and deleting p53 *in vivo*, and found that PAX7-CreER mice developed RMS and undiffereniciated pleomorphic sarcoma (UPS), whereas MyoD-CreER mice developed UPS (Blum et al., 2013). However, one caveat for the use of prior MyoD-Cre models with p53 and K-Ras alterations, which generated more undifferentiated soft tissue sarcomas, was the *ex vivo* induction of the recombination events. It has been reported that there are potential differences in myogenic promoter activity that appear to be associated with *in utero* versus neonatal and older (Rubin et al., 2011). Thus, our findings that *in utero* induction of p53 and K-Ras perturbations lead to histological and molecular signatures consistent with rhabdomyosarcoma could be attributed to this facet of developmental biology. However, additional studies are essential to further probe these features, which will help delineate the molecular facets contributing to this disease spectrum.

Interestingly, analysis of the p53-only altered tumors demonstrated signatures were more consistent with human UPS (Tsumura et al., 2006) (Doyle et al., 2010), while the knockin of the mutant K-Ras not only significantly accelerates tumor formation, but also drives the molecular signatures that highly correspond to human fusion positive and negative RMS. Therefore, our models can potentially be used to gain additional insights into the critical molecular signatures differentiating this spectrum of soft tissue sarcomas.

Besides providing valuable models to gain insights into the molecular events that drive rhabdomyosarcoma initiation and progression, our GEMM is extremely valuable secondary to the tumor-derived cell lines. As we have preliminarily demonstrated a significant advantage of these murine RMS cell lines is that they allow for highly efficient syngeneic orthotopic tumor development in immunocompetent mice. Furthermore, upon venous injection, these cell lines are also capable of rapidly establishing metastatic lung disease, thus providing additional resources for not only the evaluation of molecular and pharmacological targeting of primary, but also metastatic disease. Finally, we provide additional evidence that the murine RMS cell lines can be surrogate, and highly complementary models for evaluation of targeted and chemotherapy secondary to their very comparable responsiveness to human cell lines for various therapies, including mimicking enhanced sensitivity of the Ras mutant RD human cell line to the MEK inhibitor, Trametinib.

Ongoing and future directions include the utilization of these models to further dissect molecular mechanisms of rhabdomyosarcoma metastasis, characterization of the primary and metastatic tumor immune microenvironment and *in vivo* studies further probing identified metastatic signatures and small molecules not only targeting critical tumor intrinsic signaling pathways, but also immunomodulatory agents using these immunocompetent models.

In conclusion, we have created a novel GEMM of RMS with metastatic capabilities in addition to the derivation of tumor model resources that can be utilized for further molecular and therapeutic studies necessary to developing more effective treatment regimens for high-risk RMS patients.

## Materials and Methods

### Cell lines

Human RMS cell lines (RD and Rh30) were purchased from ATCC (Manassas, VA, USA). The cells were grown and maintained in RPMI-1640 medium (Thermo Fisher Scientific, Waltham, MA, USA) supplemented with 10% fetal bovine serum (FBS; Thermo Fisher Scientific), 1% penicillin/streptomycin (Thermo Fisher Scientific), and cultured in a humidified 5% CO_2_ incubator at 37 °C.

### Mice

Transgenic mouse lines were generated through MyoD promoter-driven Cre recombinase with core enhancer, which directs the early activation of MyoD, as reported by Chen et al (Chen et al., 2005). These mice were backcrossed using the C57BL/6J strain and further crossed to either floxed p53 mice, LSL-p53R172H, or K-Ras-G12D mice. Genotyping was performed using PCR. All research was performed in compliance with the BCM IACUC (Baylor College of Medicine, Animal Protocol AN-5225) and AAALAC recommendations as published in The Guide for the Care and Use of Laboratory Animals (NRC1996). All animals underwent comprehensive necropsies following humane euthanasia, and all major organs and tumors were evaluated. Tissues suspected of metastatic disease were submitted for histological analysis.

### Murine cell lines

Murine RMS cell lines (KMR19, KMR46, KMR72 and KMR78) were isolated from primary K-Ras-mutated RMS tumors. Resected tumors were diced and dissociated using a Tumor Dissociation Kit, mouse (Miltenyi Biotec, Cologne, Germany) and gentleMACS Dissociator (Miltenyi Biotec). The dissociated cells were resuspended and filtered through a 70μm cell strainer (Corning), then seeded, established, and sub-cultured.

### Tumor injection

KMR19, KMR46, KMR72, KMR78, RD, Rh30, and PDX40 cells were harvested and resuspended at a concentration of 1×10^6^ cells per 25 μls of PBS and then mixed with 25 μl of Matrigel matrix (Corning, Corning, NY, USA). The cell-Matrigel mixture was subsequently injected into the gastrocnemius of C57BL/6J mice, and the tumors were resected once palpable.

KMR19, KMR46, KMR72 and KMR78 were harvested and resuspended at a concentration of 1×10^6^ cells per 200 μl of PBS. The cells were injected into the tail vein of the mice, which were subsequently sacrificed a week after injection.

### Cell viability assay

To assess cell viability, cells were seeded at 5×10^3^ cells per 96-well plate (Corning, Corning, NY, USA) for 24hours and then incubated with various concentrations of vincristine, irinotecan, SN38, MK1775, and Trametinib (Selleck Chemicals, Houston, TX, USA) for 72 hours. Subsequently, the degree of cell viability was evaluated using the Cell Counting Kit-8 (Dojindo, Kumamoto, Japan) according to the manufacturer’s protocol.

### Immunohistochemistry

GEMM and orthotopic model-derived xenograft tumors, as well as clinical specimens, were embedded in paraffin, fixed, and analyzed by hematoxylin, eosin, myogenin (1:300) (EPR4789, polyclonal, Abcam, Cambridge, UK), and desmin (1:300) (PA5-16705, polyclonal, Waltham, MA, USA) staining. For IHC, VECTASTAIN ABC Kit, Rabbit IgG (Vector Laboratories, CA, USA) was used, and the stainings were visualized using Vector NovaRED Peroxidase Substrate Kit (Vector Laboratories).

### PTAH staining

For PTAH staining, we used Rapid PTAH Stain Kit (Polysciences, Warrington, PA, USA). Briefly, tumor sections slides were placed in Langeron’s iodine after deparaffinization and rehydration, and then transferred to 5% sodium thiosulfate. After rinsing in water, slides were stained in pre-heated phosphotungstic acid hematoxylin.

### Imaging

The pictures of H&E and IHC staining were got by BX53 Biological Microscope (Olympus, Tokyo, Japan). The whole pictures of lung sections were obtained by tile scanning, which merged 20x images and allowed the acquisition of overview images of the specimen, and Leica DMi8 and LAS X software (Leica Microsystems, Wetzlar, Germany) were used for the scan.

### RNA-sequencing Library preparation and Sequencing

RNA samples underwent quality control assessment using the RNA tape on Tapestation 4200 (Agilent) and were quantified using a Qubit Fluorometer (Thermo Fisher). The RNA libraries were prepared and sequenced at the University of Houston Seq-N-Edit Core per standard protocols. RNA libraries were prepared with QIAseq Stranded Total RNA Library Kit (Qiagen) using 500 ng input RNA. mRNA was enriched with Oligo-dT probes attached to Pure mRNA beads (Qiagen). RNA was fragmented, reverse transcribed into cDNA, and ligated with Illumina sequencing adaptors. The size selection for libraries was analyzed using the DNA 1000 tape Tapestation 4200 (Agilent). The prepared libraries were pooled and sequenced using NextSeq 500 (Illumina); generating ~17million 2×76 bp paired-end reads per samples.

### Total RNA-Seq analysis

Paired-end sequencing reads were trimmed using the trimGalore software. The trimmed reads were mapped using STAR against the murine genome build UCSC mm10, and quantified using featureCounts (37) against the Gencode gene model. Human RMS gene signatures were derived using the RNAseq counts data for GSE108022 which includes 101 rhabdomyosarcoma (35 fusion-positive and 66 fusion-negative) samples and 5 normal muscle. Differential expression (DE) analysis was performed using DESeq2 R package (1.28.1) (Wang et al. 2009). The p-values were adjusted with Benjamini and Hochberg’s approach for controlling the false discovery rate. Significantly differentiated genes between the comparisons were identified by applying the criteria of adjusted p-value< 0.05 and fold change exceeding 1.5x. Pathway enrichment analysis was carried out using the GSEA (http://software.broadinstitute.org/gsea/index.jsp) software package; significance was achieved for adjusted q-value<0.25.

Signature correlations were performed using normalized expression data in the Schafer (n=30), Davicioni (n=147), and Barr (GSE66533, n=58) datasets in R2:Genomics Analysis and Visualization Platform (http://r2.amc.nl). The UPS cohort RNAseq counts data was kindly provided by Dr. Nischalan Pillay (Steele et al, Cancer Cell 2019) and consisted of 58 tumor samples and 5 normal adjacent tissue controls. For each cohort a z-score was computed for each gene in a given gene signature per patient sample resulting in a patient activity score. For each combination of signatures the individual patient activity scores were plotted on the x and y axes and the Pearson Correlation Coefficient along with p-value were calculated.

## Acknowledgements

We would like to acknowledge the Advanced Technologies Core services at the Baylor College of Medicine.; CPRIT Core Facility Support Award (CPRIT-RP180672), the NIH (CA125123 and RR024574 to the Cytometry and Cell Sorting Core at Baylor College of Medicine).

## Competing Interests

The authors declare no competing interests.

## Funding

This work was supported by the National Institutes of Health (R21 CA234665) and The Faris D. Virani Ewing Sarcoma Center to JTY. C. Coarfa was partially supported by (CPRIT) RP170005, NIH P30 shared resource grant CA125123, and NIEHS P30 Center grant 1P30ES030285.

## Author contributions

KN and JTY conceived and designed the study. GEMM mice were maintained by LK. Tumor cell injections and tumor collections were performed by KN and LK. Staining of tumor samples were performed KN and RS. RNA sequencing data analysis was performed by TDP. The diagnosis of tumor was performed by BWS, and BWS, EP and CC provided study supervision. JTY provided direct supervision. KN and JTY wrote the manuscript. All authors read and approved the final manuscript.

## Notes

### Competing Interest Statement

The authors have declared no competing interest.

## References

Blum, J. M., Ano, L., Li, Z., Van Mater, D., Bennett, B. D., Sachdeva, M., Lagutina, I., Zhang, M., Mito, J. K., Dodd, L. G. et al. (2013). Distinct and overlapping sarcoma subtypes initiated from muscle stem and progenitor cells. Cell Rep 5, 933–40.

Chen, J. C., Mortimer, J., Marley, J. and Goldhamer, D. J. (2005). MyoD-cre transgenic mice: a model for conditional mutagenesis and lineage tracing of skeletal muscle. Genesis 41, 116–21.

Doyle, B., Morton, J. P., Delaney, D. W., Ridgway, R. A., Wilkins, J. A. and Sansom, O. J. (2010). p53 mutation and loss have different effects on tumourigenesis in a novel mouse model of pleomorphic rhabdomyosarcoma. J Pathol 222, 129–37.

Felix, C. A., Kappel, C. C., Mitsudomi, T., Nau, M. M., Tsokos, M., Crouch, G. D., Nisen, P. D., Winick, N. J. and Helman, L. J. (1992). Frequency and diversity of p53 mutations in childhood rhabdomyosarcoma. Cancer Res 52, 2243–7.

Hettmer, S., Liu, J., Miller, C. M., Lindsay, M. C., Sparks, C. A., Guertin, D. A., Bronson, R. T., Langenau, D. M. and Wagers, A. J. (2011). Sarcomas induced in discrete subsets of prospectively isolated skeletal muscle cells. Proc Natl Acad Sci U S A 108, 20002–7.

Ku, B. M., Bae, Y. H., Koh, J., Sun, J. M., Lee, S. H., Ahn, J. S., Park, K. and Ahn, M. J. (2017). Mutational status of TP53 defines the efficacy of Wee1 inhibitor AZD1775 in KRAS-mutant non-small cell lung cancer. Oncotarget 8, 67526–67537.

Leuschner, I., Langhans, I., Schmitz, R., Harms, D., Mattke, A., Treuner, J., Kiel Pediatric Tumor, R. and the German Cooperative Soft Tissue Sarcoma, S. (2003). p53 and mdm-2 expression in Rhabdomyosarcoma of childhood and adolescence: clinicopathologic study by the Kiel Pediatric Tumor Registry and the German Cooperative Soft Tissue Sarcoma Study. Pediatr Dev Pathol 6, 128–36.

Marampon, F., Ciccarelli, C. and Zani, B. M. (2006). Down-regulation of c-Myc following MEK/ERK inhibition halts the expression of malignant phenotype in rhabdomyosarcoma and in non muscle-derived human tumors. Mol Cancer 5, 31.

McKinnon, T., Venier, R., Dickson, B. C., Kabaroff, L., Alkema, M., Chen, L., Shern, J. F., Yohe, M. E., Khan, J. and Gladdy, R. A. (2015). Kras activation in p53-deficient myoblasts results in high-grade sarcoma formation with impaired myogenic differentiation. Oncotarget 6, 14220–32.

Rubin, B. P., Nishijo, K., Chen, H. I., Yi, X., Schuetze, D. P., Pal, R., Prajapati, S. I., Abraham, J., Arenkiel, B. R., Chen, Q. R. et al. (2011). Evidence for an unanticipated relationship between undifferentiated pleomorphic sarcoma and embryonal rhabdomyosarcoma. Cancer Cell 19, 177–91.

Stewart, E., Federico, S. M., Chen, X., Shelat, A. A., Bradley, C., Gordon, B., Karlstrom, A., Twarog, N. R., Clay, M. R., Bahrami, A. et al. (2017). Orthotopic patient-derived xenografts of paediatric solid tumours. Nature 549, 96–100.

Stratton, M. R., Fisher, C., Gusterson, B. A. and Cooper, C. S. (1989). Detection of point mutations in N-ras and K-ras genes of human embryonal rhabdomyosarcomas using oligonucleotide probes and the polymerase chain reaction. Cancer Res 49, 6324–7.

Subramanian, A., Narayan, R., Corsello, S. M., Peck, D. D., Natoli, T. E., Lu, X., Gould, J., Davis, J. F., Tubelli, A. A., Asiedu, J. K. et al. (2017). A Next Generation Connectivity Map: L1000 Platform and the First 1,000,000 Profiles. Cell 171, 1437–1452 e17.

Tsumura, H., Yoshida, T., Saito, H., Imanaka-Yoshida, K. and Suzuki, N. (2006). Cooperation of oncogenic K-ras and p53 deficiency in pleomorphic rhabdomyosarcoma development in adult mice. Oncogene 25, 7673–9.

Wang, C. (2012). Childhood rhabdomyosarcoma: recent advances and prospective views. J Dent Res 91, 341–50.

Wilke, W., Maillet, M. and Robinson, R. (1993). H-ras-1 point mutations in soft tissue sarcomas. Mod Pathol 6, 129–32.

Yohe, M. E., Gryder, B. E., Shern, J. F., Song, Y. K., Chou, H. C., Sindiri, S., Mendoza, A., Patidar, R., Zhang, X., Guha, R. et al. (2018). MEK inhibition induces MYOG and remodels super-enhancers in RAS-driven rhabdomyosarcoma. Sci Transl Med 10.

Yohe, M. E., Heske, C. M., Stewart, E., Adamson, P. C., Ahmed, N., Antonescu, C. R., Chen, E., Collins, N., Ehrlich, A., Galindo, R. L. et al. (2019). Insights into pediatric rhabdomyosarcoma research: Challenges and goals. Pediatr Blood Cancer 66, e27869.

