## Supplemental Figure 1 for "MyoD-Cre driven alterations in K-Ras and p53 lead to a mouse model with histological and molecular characteristics of human rhabdomyosarcoma with direct translational applications"

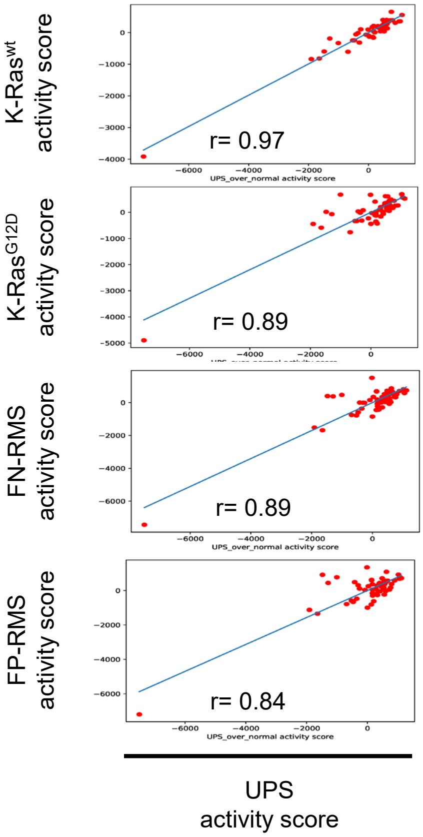


**Supplemental Figure 1. The correlation of RMS GEMM tumors and human FN and FP-RMS to human Undifferentiated Pleiomorphic Sarcoma**

Pearson correlation coefficients for signature correlations using UPS DEGs are shown for murine $Ras^{WT}$ GEMM tumors, K-$Ras^{G12D}$ GEMM knockin tumors, and human FN and FP signatures.
